# RELATIONSHIP BETWEEN ACTUAL STRESS AND IN SITU MORPHOLOGICAL CHANGES IN THE MECHANICAL BEHAVIOR OF BIOMIMETIC POROUS HIERARCHICAL SCAFFOLDS

**DOI:** 10.1101/2024.12.27.630537

**Authors:** Gregorio Marchiori, Nicola Sancisi, Gianluca Tozzi, Massimiliano Zingales, Gaia Prezioso, Andrea Visani, Andrea Zucchelli, Alberto Sensini

## Abstract

This study investigates the evolution with strain of the material volume fraction (i.e., porosity) and geometry in porous scaffolds to obtain a more accurate description of their stress-strain behavior.

Single bundles and hierarchical structures (8 bundles enveloped by a membrane) were produced by electrospinning as tendon/ligament scaffolds. They underwent a micro-tomography *in situ* tensile test. Apparent and net stress were obtained using the initial sample cross-section and material volume fraction to normalize axial force. Micro-tomography revealed sample morphology change with strain to calculate the actual stress-strain. Moreover, nanofibers arrangement was revealed by scanning electron microscopy on both bundles and membranes.

The description of the mechanical response significantly changed using evolving morphometry (actual stress-strain) instead of initial static one (apparent stress-strain), for both single bundle and hierarchical structure. The actual elastic modulus of the single bundles (583±97 MPa) was statistically higher than that of the hierarchical structures (163±107 MPa). This is related to the membrane, membrane-bundle and inter-bundle interactions. In the hierarchical structure, portions of the material resisting traction are constituted by nanofibers not aligned with the load. The different definitions for the stress-strain behavior allow different accuracy levels depending on the experimental complexity.

The evolution of morphology with deformation can significantly affect the description of the mechanical response of porous scaffolds. This has a double impact in practical applications: at the body scale, it allows a better comparison between the scaffold behavior and the target tissue; at the cellular scale, it predicts the actual substrate stiffness that cells will face.

**Graphical Abstract:** 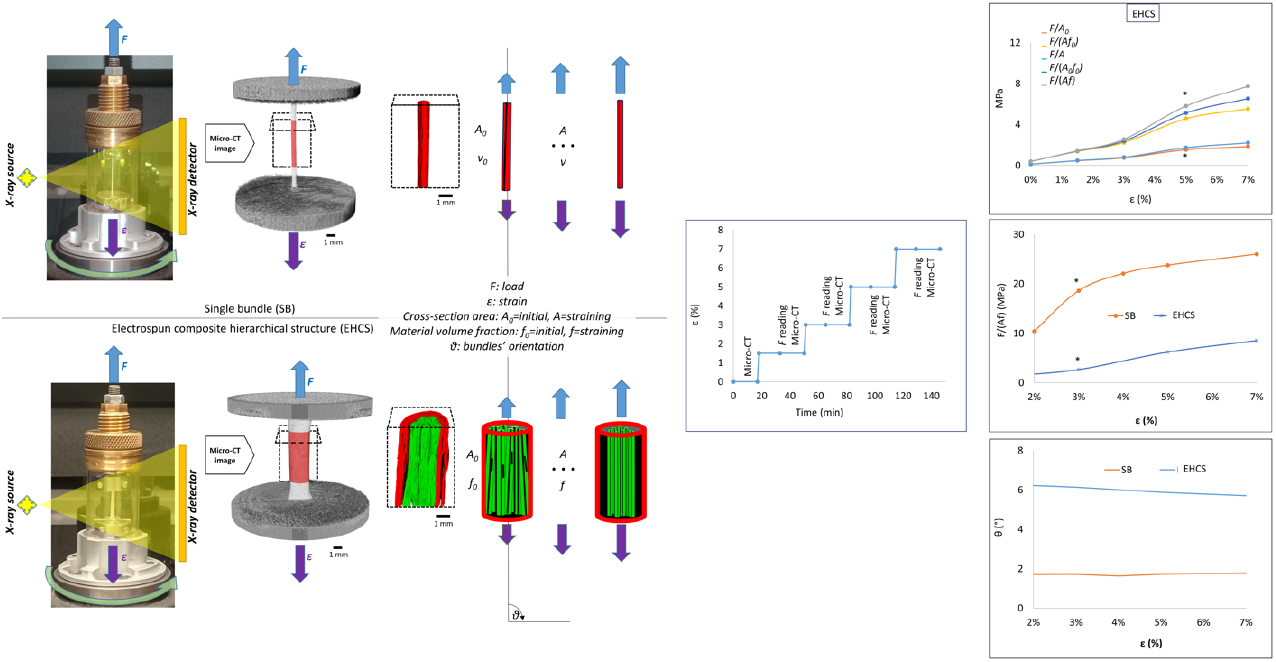

## 1. Introduction

The hierarchical structure of load-bearing tissues is optimized for their specific biomechanical function and anatomical region [1]. This is the case of tendons and ligaments (T/L), in both cases constituted by dense hierarchical fibrous collagenous tissue [2], whose fibers are primarily able to resist tensile loading [3]. However, there are peculiar differences between tendons and ligaments [4], considering their different role: bone-bone connection for ligaments and muscle-bone connection for tendons. Morphological differences were also reported between tendons [5] when harvested from different anatomical locations.

A significant clinical scenario in this context is the rupture of T/L, a common debilitating orthopedic injury. Damaged tendons – such as Achilles, patellar and quadriceps – and ligaments – such as anterior cruciate, posterior cruciate and collaterals – may need reconstruction by a graft. Standard procedures use biological grafts, i.e. autografts or allografts. However, they do not represent the optimal solution in many cases, such as revision after graft failure, risk of rejection for allografts, need for rapid recovery. Therefore, synthetic grafts had been considered starting from the eighties, but they soon declined because of low results in the medium follow-up and common secondary inflammations. Then, innovative artificial grafts became a valid solution for very specific cases, such as anterior cruciate ligament reconstruction in subjects with more than 40 years and the need for a rapid recovery [6], multi-ligamentous lesions, and tissue reinforcement. Various synthetic grafts exist but they still often have unsatisfactory outcomes at both the short and long term of clinical applications [7]. However, the market of artificial T/L is expected to strongly increase globally [https://www.exactitudeconsultancy.com/reports/28949/artificial-tendons-and-ligaments-market]. For commercial success, the target product should be biocompatible, functional, and tunable for many clinical cases. Thus, scaffolds for T/L tissue engineering should be customized to replicate the hierarchical structure and the biomechanical properties of the various T/L of interest. Among the different biofabrication techniques, electrospinning emerged as one of the most promising ones to produce biomimetic T/L scaffolds, replicating the T/L structure from the fascicle [8][9][10][11] up to the whole tissue level [12]. These scaffolds could also modify fibroblast morphology and orientation in static and dynamic conditions, demonstrating suitability for *in vivo* studies [13].

When targeting the material’s mechanical characteristics, attention should be paid to their definition. In the study of Woo et al. [14], the effects of donor age and specimen orientation on the tensile properties of the femur-ligament-tibia complex for anterior cruciate ligament (ACL) were analyzed by reading load (N) and elongation (mm) and comparing stiffness (N/mm), ultimate load (N) and absorbed energy (Nm). These characteristics are named “extrinsic”, being proper of the tested complex and not of the specific tissue. A step towards tissue’s “intrinsic” (i.e., constitutive) properties comes from measuring the specimen’s initial cross-section (mm^2^) and length (mm) to transform force-displacement characteristics into engineering (i.e., apparent) stress (N/mm^2^ = MPa)-strain (mm/mm). This was used, for instance, to compare the mechanical characteristics of the patellar tendon and knee ligaments [15][16]. Engineering stress was also identified as a good metric for optimal pre-tensioning of ACL grafts [17] and to show how a functional scaffold improves the biomechanical properties of regenerated tendons [18]. It also allows a direct comparison of the scaffolds’ mechanical performances with the properties of the target tissue, but causes a relevant underestimation of the scaffolds’ actual mechanics [18]. This is mainly due to void spaces in porous structures, causing a dramatic underestimation of the actual stress supported by their cross-section.

Due to their simplicity, engineering stress-strain characteristics have become a standard: few geometrical measures on the specimen are needed, taken at the beginning of the mechanical test using a caliper or a photo with scale. However, they may not describe the actual constitutive state of the material due to: (i) section changes during the mechanical test (i.e., with progressing deformation); (ii) presence of voids (i.e., specimen density is not equal to material density); (iii) non-homogenous strain, e.g. it does differ from clamped to middle portions. To consider points (i) and (ii), the evolution of micro-structure with deformation in the ACL was analyzed [19]. Point (ii) was also considered for electrospun T/L scaffolds, by calculating the initial material volumetric fraction to transform apparent stress in net stress [20][21][12][22]. Point (iii) was considered by applying digital volume correlation (DVC) on electrospun scaffolds and materials in general [20].

This study has the scope of advancing the mechanical description of tissue engineering poly-L(lactic) acid and collagen type I composite hierarchical scaffolds by comparing the impact of different evaluation methods and further extending the metrics to characterize the scaffolds’ mechanical characteristics. The effectiveness of considering the net stress for porous scaffolds is determined. In addition, the morphological quantities that define the mechanical characteristics are used both in a static (i.e., determined before a test and then unchanged) and dynamic (i.e., measured and updated at every step of the tensioning test) way. A micro-CT *in situ* test was implemented to measure the change with deformation of specimens’ section and material volumetric fraction and to quantify the effect on stress-strain characteristics of net vs apparent stress and of updated vs not updated morphological quantities. Moreover, the same test contributed to relating mechanical response peculiarities and morphological changes of scaffolds during the application of loads.

Therefore, the study aims to change the perspective in characterizing the mechanical behavior of tissue scaffolds, specifically by investigating (i) with a non-standard, complex, experimental testing, the hypothesis that strain-dependent morphology affects the stress metrics; (ii) how to upgrade standard metrics and to implement morphological strain-dependency in a standard testing set-up integrated with few, simple solutions.

As a whole, the results will give important general indications for the actual mechanical description and design of hierarchical, porous materials, e.g., of advanced scaffolds for tissue engineering.

## 2. Materials and Methods

### 2.1 Materials

Bovine skin’s collagen type I (Coll) (Kensey Nash Corporation DSM Biomedical, Exton, USA) and poly-L(lactic) acid (PLLA) (Lacea H.100-E, Mw = 8.4 × 104 g mol^−1^, PDI = 1.7, Mitsui Fine Chemicals, Dusseldorf, Germany) were used to prepare the scaffolds. As a solvent system, a mix of 2,2,2-trifluoroethanol (TFE), 1,1,1,3,3,3-Hexafluoro-2-propanol (HFIP) (Sigma-Aldrich, Staint Louis, USA) was used in a percentage of 50:50 (v/v). As crosslinkers for collagen, N-(3-Dimethylaminopropyl)-N′-ethylcarbodiimide hydrochloride (EDC), N-hydroxysuccinimide (NHS) and 95% ethanol (Sigma-Aldrich, Staint Louis, USA) were employed.

### 2.2 Electrospun scaffolds preparation

A polymeric blend of PLLA/Coll-75/25 (w/w) obtained from a 18% (w/v) solution of PLLA and Coll dissolved in TFE:HFIP = 50:50 (v/v) was employed to electrospun nanofibers mimicking the morphology of T/L fibrils [23][24][25]. Electrospun T/L fascicle-inspired bundles of aligned nanofibers were produced following a consolidated method [26][27][28]. To reach a diameter in the range of human T/L fascicles (500-650 μm), an electrospinning machine (Spinbow srl, Bologna, Italy) with a high-speed rotating drum collector (length = 405 mm, diameter = 150 mm; peripheral speed =19.6 m s^−1^; drum rotations = 2,500 rpm) was adopted. To facilitate the detachment of the nanofibers’ mats, the drum was coated with a sheet of polyethylene paper (Turconi S.p.A, Italy). The following electrospinning parameters were used: an applied voltage of 22 kV; a feed rate of 0.5 mL h^−1^ (syringe pump KD Scientific 200 series, IL, United States) used to deliver the solution by four metallic needles (internal diameter = 0.51 mm, Hamilton, Romania) connected to the syringes by polytetrafluoroethylene tubes (Bola, Germany); needles-collector distance was 200 mm; 180 mm of excursion of the sliding spinneret supporting the needles, with a speed of 1,500 mm min^−1^; 2h of electrospinning session; room temperature and relative humidity of 20–30%. At the end of the electrospinning session, the mat was cut in circumferential stripes of 45 mm, rolled up and pulled off the drum obtaining single bundles (SB) of aligned nanofibers (Fig. 1b-c). To simulate the whole hierarchical structure of T/L [23][24][25] (Fig. 1a), electrospun hierarchical composite scaffolds (EHCS) (Fig. 1d-e) were assembled adapting a consolidated procedure [21][13][29]. N=8 SB per each EHCS were grouped and covered with an electrospun epitenon/epiligament-inspired membrane. For this procedure, a second electrospinning machine (Spinbow srl, Bologna, Italy) using a high-voltage power supply (FuG Elektronik GmbH, Schechen, Germany), and a syringe pump (KD Scientific Legato 100, Illinois, USA) was used. Each assembly was placed in a custom-made machine equipped with a flat plate aluminum collector, rotating the group of bundles during the electrospinning session. To produce the membrane, the group of bundles was maintained in a static position alternated by rotation sessions (5 sessions of 10 rpm for 30 sec every 20 min of stasis) (Fig. 1d). The PLLA/Coll-75/25 solution and the electrospinning parameters were the same as previously described. SB and EHCS were finally crosslinked with a crosslinking solution of EDC and NHS 0.02 M in 95% ethanol, following a consolidated procedure [27]. After crosslinking, SB and EHCS were cut in segments of 20 mm for the following characterizations.

**Figure 1.**
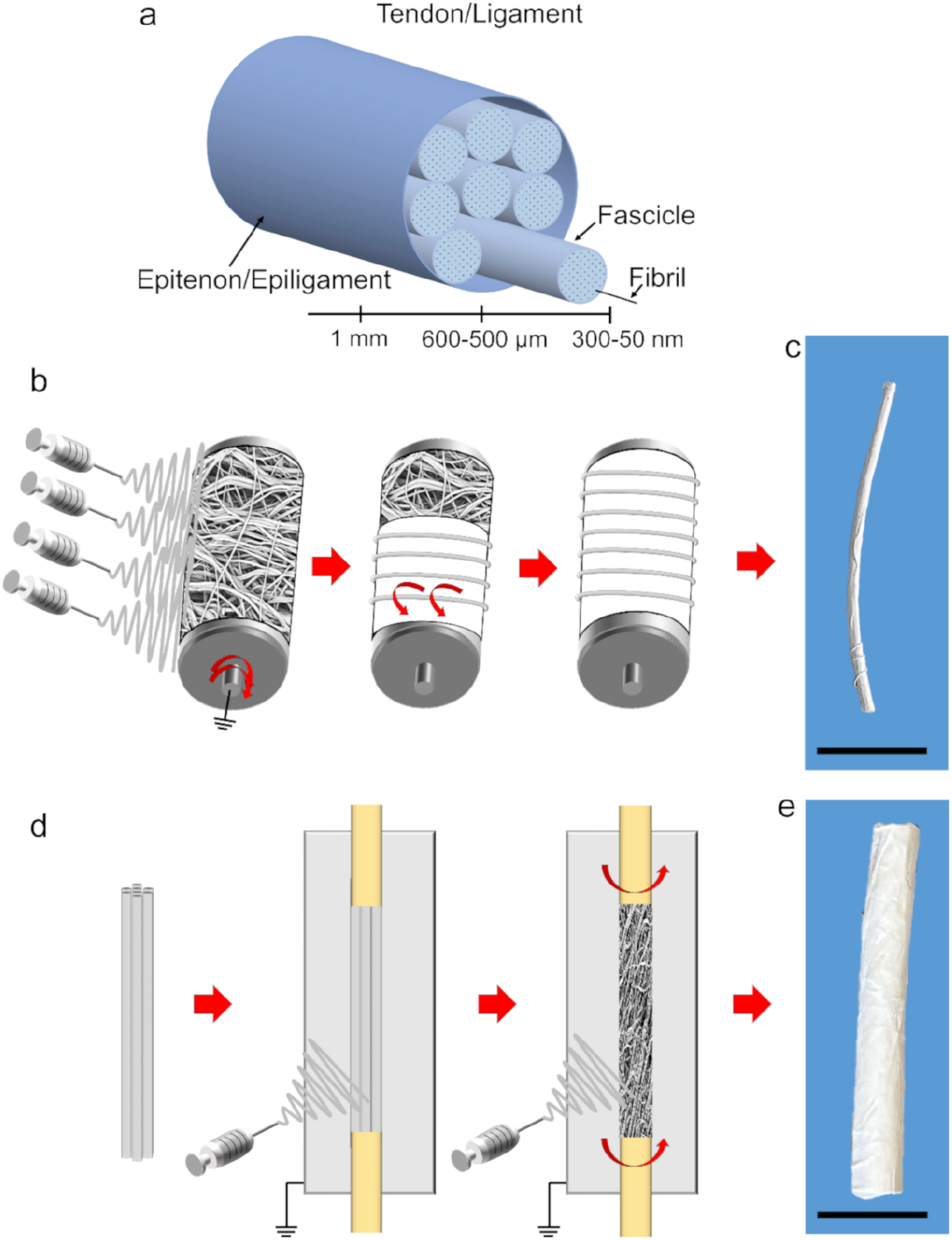
Structure of natural tissues and scaffolds’ fabrication. A) Hierarchical structure of T/L; b) electrospun SB fabrication; c) typical SB (scale bar = 5 mm); d) EHCS production; e) typical EHCS (scale bar = 5 mm).

### 2.3 SEM

To investigate the SB and EHCS morphology, a Scanning Electron Microscope (SEM) (Phenom Pro-X, PhenomWorld, Eindhoven, Netherlands) was used. Samples were gold-sputtered and then imaged at 10 kV. The mean and standard deviation (SD) of the nanofiber diameters of both SB and EHCS membranes (magnification = 8,000x) were obtained by measuring 100 nanofibers with ImageJ [30]. SB and EHCS diameters were measured via an optical microscope (Axioskop, Zeiss, Pleasanton, CA, United States) connected to a camera (AxioCam MRc, Zeiss, Pleasanton, CA, United States) as mean and SD of 20 measures. The nanofiber orientation was investigated with the Directionality plugin of ImageJ [31], by adopting a consolidated method [32]. In brief, the Local Gradient Orientation method of Directionality was used for both bundles and EHCS membrane on four images (magnification = 8,000x) along the scaffold’s axis. The results were reported as mean and SD between the four images, 0° corresponding to nanofibers’ axial orientation.

### 2.4 Micro-CT *in situ* test

SB (n=4) and EHCS (n=4) samples underwent micro-CT *in situ* mechanical tests. The initial length (*L*_*0*_, mm) and section (diameter *d*_*0*_ , mm; area 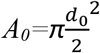, mm^2^) of the dried samples were measured using ImageJ on photographic macro images (Fig. 2a), while their mass (*w*) by a micro-balance. The initial volume fraction (*f*_*0*_) was calculated as:

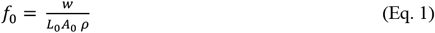

where *ρ* is the density of the blend and is equal to 1.27 g cm^-3^ [33].

**Figure 2.**
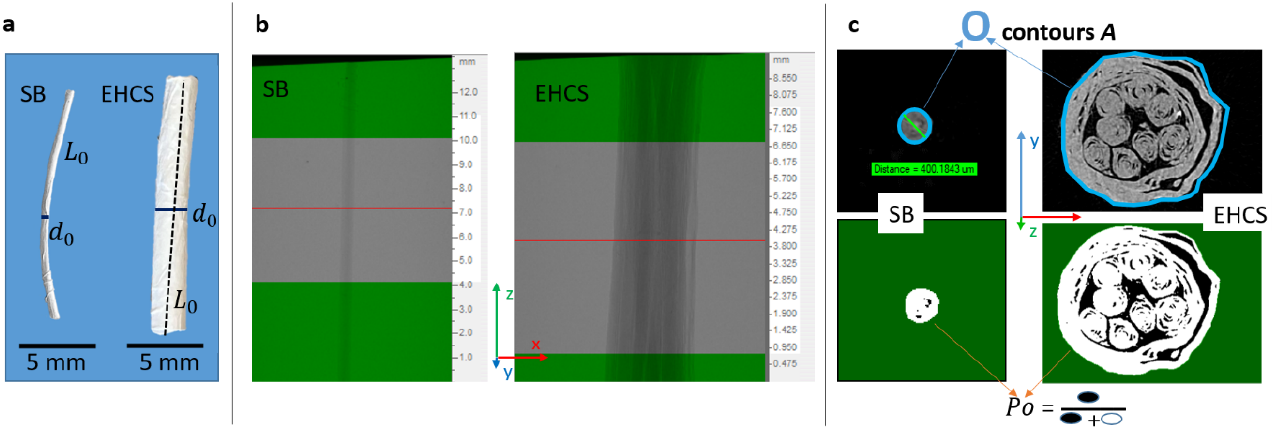
(a) Photography of SB and EHCS sample with schematic measure of diameter (d_0_) and length (L_0_), where L_0_ was obtained using a broken line in case of curvature; (b) micro-CT radiography of SB and EHCS sample highlighting the vertical (z) length of the region of interest (ROI, out-of-green), and a cross-section (x-y, red line); (c) grey-level (up) and binarized (down) micro-CT cross-section (x-y) of SB and EHCS sample highlighting ROI (out-of-green), relative actual section area (A’, contoured in blue) and porosity (Po, ratio between voids in black, and voids + material in white).

Each sample was then immersed in PBS for 2 minutes before being clamped in the micro-CT testing system (MTS *in situ* mechanical tester for Skyscan 1172, Bruker, Belgium; maximum actuator displacement = 5.5 mm, displacement rate = 0.001 mm s^-1^). The gauge length of the samples was measured as the initial clamp-clamp distance (≈ 10 mm for both SB and EHCS length, Fig. 3, Video S1, Video S2) from a single micro-CT image at the minimum strain allowed by the load cell (0.45 N, corresponding to about 2% strain for SB and 0% strain for EHCS). The first micro-CT scan was acquired under these conditions. Based on the gauge length, testing system displacements were imposed to reach defined strains for SB (i.e. 3%, 4%, 5%, 7%) and for EHCS (i.e., 1.5%, 3%, 5%, 7%); corresponding tensile forces (*F*, N) were read by the load cell (max 200 N). These specific strain levels were chosen as relevant in the force-strain curves of previous *ex situ* mechanical tests on the same kind of samples [33]. A 15-minute stress-relaxation period was applied at each strain step to stabilize the sample microstructure, followed by a micro-CT acquisition [34].

**Figure 3.**
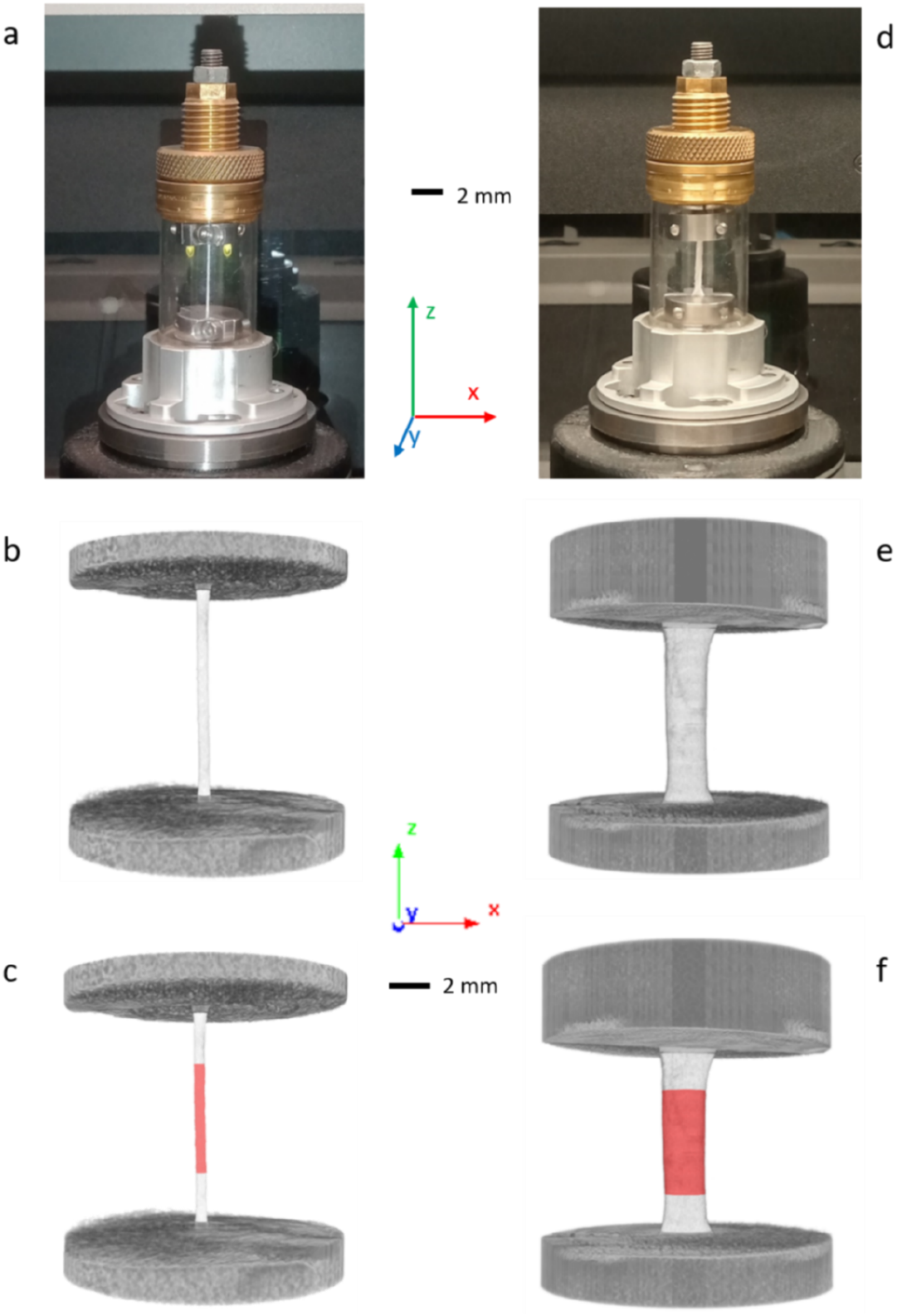
SB (left) and EHCS (right) specimen in the micro-CT in situ tester (a,d), frontally imaged with clamps only (b,e) and highlighting ROI in red (c,f). Renderings by SkyScan CT-Vox software (Supplementary videos 1 and 2).

SB were scanned with an applied voltage of 40 kV and a current of 75 µA. The scan orbit was 180° with a rotation step of 0.8°, and 4 frames were averaged for each rotation angle; the voxel size was 13 µm, resulting in a scan duration of ∼ 17 minutes. The scanning protocol for EHCS was the same, except for decreasing the voxel size to 9 µm to better discriminate between bundles and membranes. The image reconstruction was carried out with a modified Feldkamp algorithm using the SkyScanTM Nrecon software accelerated by GPU, applying a level-5 ring-artifact correction without beam-hardening correction and smoothing.

Region of interest (ROI) selection and morphometric analysis were all performed using SkyScan CT-Analyser software. ROI extended 6 mm in the vertical direction, which is 10 mm of gauge length minus 2 mm from each steel clamp to avoid metal artifacts (Fig. 2b and 3c,f), and wrapped the sample on the cross-section (Fig. 2c). In EHCS, membranes and bundles (Fig. 2 c and 4f,g) could not be separated in a fully automatic way by morphological operations, because of the tightening in traction and overlapping grey intensity distributions; therefore, a semi-automatic procedure was followed for rendering membrane and bundles separately (Fig. 4 f,g and Fig. 5 c,d). For each sample and strain level, the following parameters were calculated:

**Figure 4.**
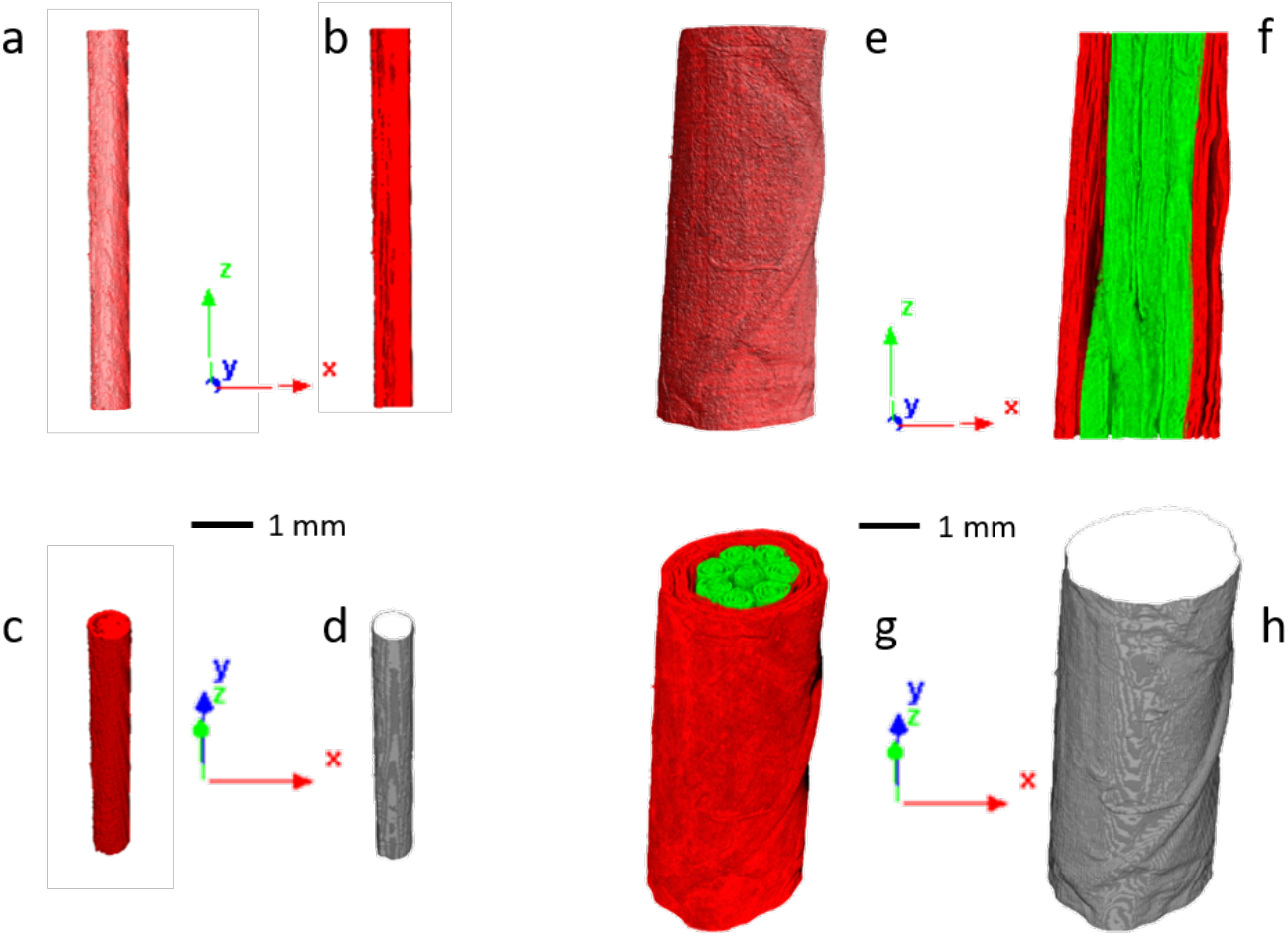
SB (left) and EHCS (right) specimen frontal view (a,e), binarized and frontally cut (b,f) or transversally cut (c,g) to highlight material and void, wrapped by ROI (d,h) to highlight transversal section. For EHCS, bundles are showed in green and membrane in red. Renderings by SkyScan CT-Vox software.

**Figure 5.**
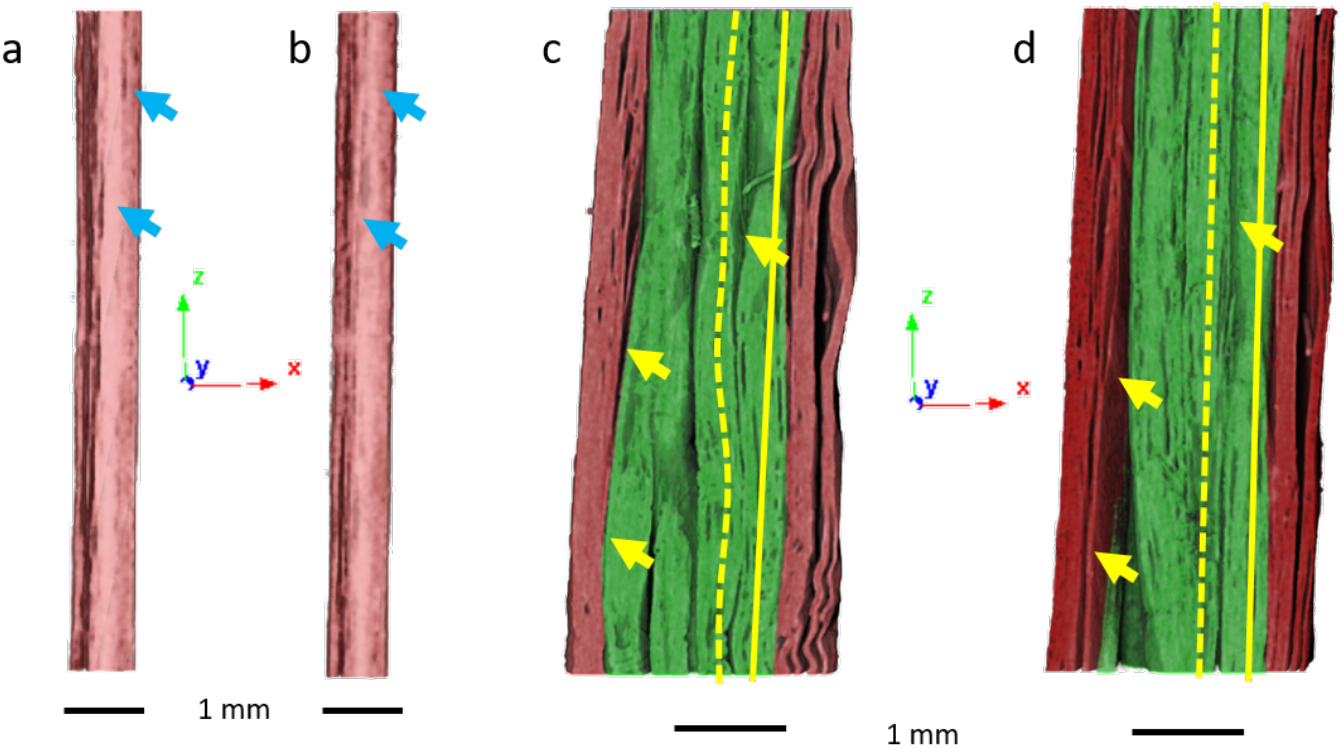
Frontal view cut of SB at 2% strain (a) and 7% strain (b), of EHCS at 0% strain (c) and 7% strain (d), highlighting changes in porosity (arrows), bundle’s tortuosity (dotted lines) and orientation (full lines).

1. Micro-CT porosity (*Po*, %), volumetric fraction (*f’*, %) and section (*A’*, mm^2^): after binarization, *Po* is the percentage of measurable void space inside the sample ROI volume (3D analysis in CT-Analyser, Fig. 2c and Fig. 4b,c,f,g) and *f’=*1*-Po*. Since part of voids cannot be measured, *f’* is significantly higher than the actual volumetric fraction. It is assumed that the percentage of unmeasured voids remains constant, so the ratio *k*=*f*_0_/*f*′_0_ is assumed constant, where *f*_0_ is the volume fraction measured at Eq. 1, and *f*_*0*_*’* is the volume fraction obtained from *Po* at strain 0% (the first micro-CT scan for EHCS, extrapolated from 2%-7% strain trend for SB). *A’* is the ROI cross-section area averaged on the 451 transversal sections along the ROI vertical length (2D analysis in CT-Analyser, Fig. 2c and Fig. 4d,h). In analogy with volumetric fraction, *y*=*A*_0_/*A*′_0_ is assumed constant.
2. Actual volumetric fraction (*f*): volumetric fraction considering the unmeasured voids (Fig. 5a,b,c,d):

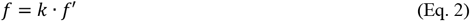
3. Actual section (*A*):

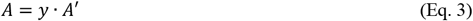
4. Apparent stress (MPa):

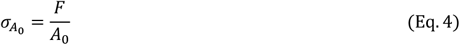
5. Apparent net stress (MPa):

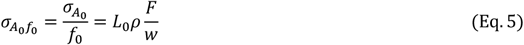
6. Actual stress (MPa):

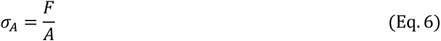
7. Net stress (MPa):

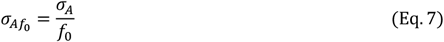
8. Actual net stress (MPa):

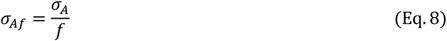
9. Estimated net stress (MPa): to predict cross-section area change, the first- and second-order volume change of the specimen was considered by Poisson’s ratio *ν*:

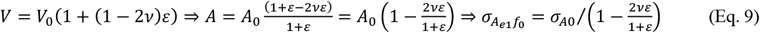

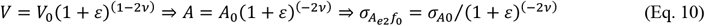
10. Bundle orientation (*θ*, °): bundle average angle with respect to the loading axis, i.e., the angle between the loading axis and the line connecting the top and bottom centroids of the bundle (0° = parallel, details in [33]). It is a single value in the case of SB, while it is the mean of the 8 bundles in EHCS (Fig. 5c,d).
11. Bundle tortuosity (*τ*): bundle length divided by the ROI vertical length, where bundle length is automatically calculated by the CT-Analyser software as the length of a rod equivalent to the bundle in terms of volume and diameter; briefly, the software calculates the average diameter of an object (i.e., bundle) and its volume, considers a rod with those geometrical values and derives its length; *τ* is 1 when the bundle is perfectly aligned with the loading axis and without crimps (details in [33]). It is a single value in the case of SB, while it is the mean of the 8 bundles in EHCS (Fig. 5c,d).
12. Stress-strain gradient: d(·)/d*ε*, where *ε* is strain as absolute value and (·) is 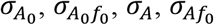 or σ_*Af*_, in MPa.

## 3. Results and discussion

After the fabrication process, nanofibers, SB and EHCS showed a biomimetic morphology that closely resembles the structure of fibrils, fascicles, membranes and whole T/L. The overall results confirmed previous analyses performed by some of the present authors [20], thus are only summarized here. Specifically, SB (cross-sectional diameter = 0.6 ± 0.1 mm; weight = 2.0 ± 0.3 mg, *f*_*0*_ = 0.3 ± 0.1) and EHCS (cross-sectional diameter = 2.1 ± 0.1 mm; weight = 29.0 ± 4.4 mg, *f*_*0*_ = 0.3 ± 0.1) displayed morphology (Fig. 1c,e and 6a,d) and thickness of natural T/L reported in the literature [23][2][24][25]. The SEM investigation showed that nanofibers of bundles (diameter = 230 ± 60 nm) (Fig. 6c,e) and membranes (diameter = 262 ± 80 nm) (Fig. 6d,e) were continuous and without beads, with the same order of magnitude as T/L collagen fibrils [23][24][25]. Differences between bundle and membrane nanofibre diameters were due to the different fabrication procedures (Fig. 6e). The orientation analysis confirmed a preferential axial distribution for bundles’ nanofibers and a circumferential orientation for membranes, though with a more random distribution, similar to our previous studies (Fig. 6f) [13,20]. This analysis confirmed the morphological biomimicry of PLLA/Coll nanofibers with the ones of T/L fascicles and their membranes [23][24][25]. Even if the nanofibers of bundles and membranes were completely the same in terms of composition, the different electrospinning strategies contributed to modifying their orientation and diameter distribution while remaining closely similar to their biological counterparts. The stress-strain differences between SB and EHCS thus depend on the membrane’s nanostructure, and on the inter-bundle and membrane-bundle interaction in EHCS.

**Figure 6.**
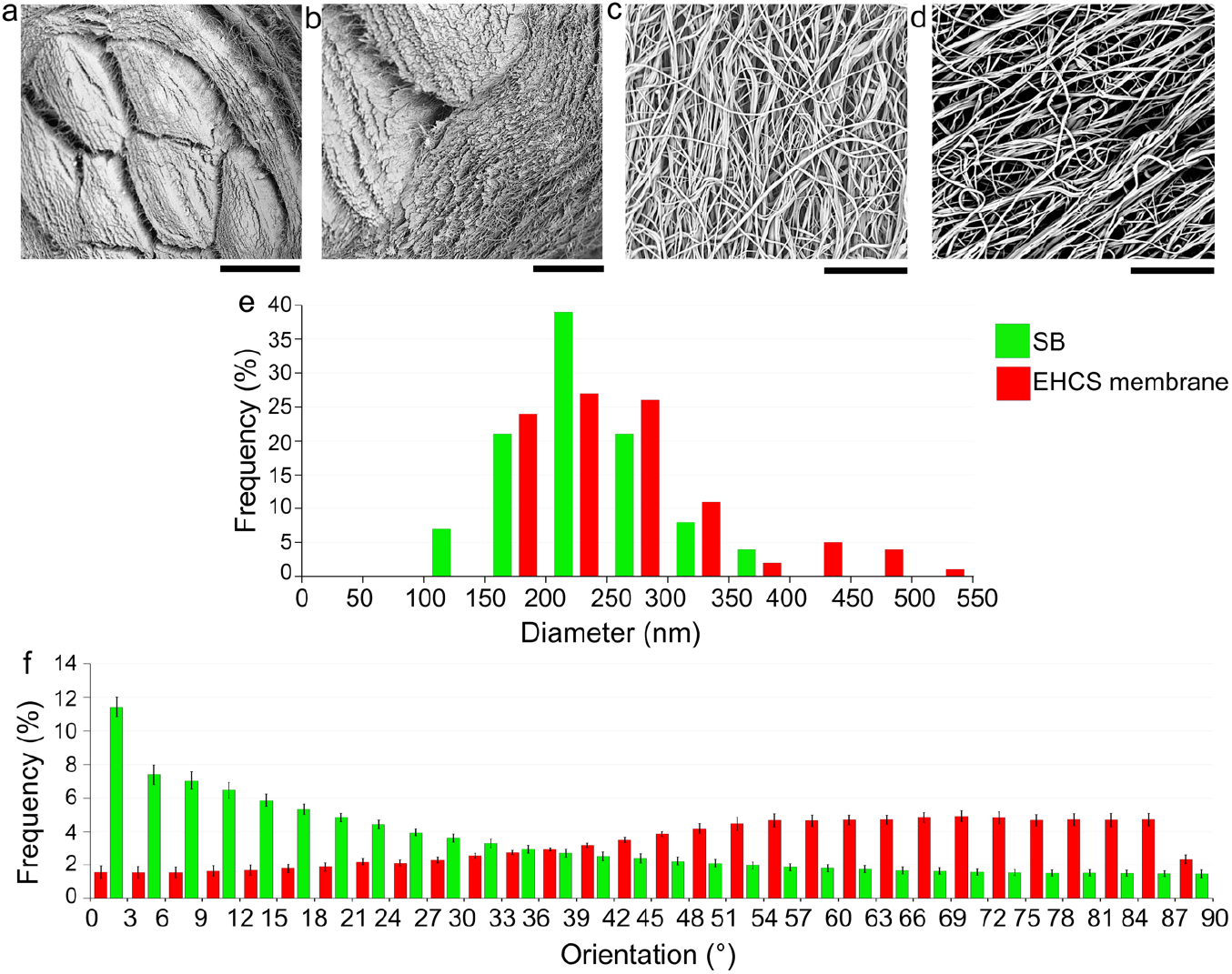
SEM investigation of scaffolds: a) overview of EHCS with its 8 SB inside (scale bar = 300 μm; magnification = 250x ); b) detail of internal SB and membrane (scale bar = 100 μm; magnification = 680x ); c) SB nanofibers (scale bar = 10 μm; magnification = 8000x); d) nanofibers of EHCS membrane (scale bar = 10 μm; magnification = 8000x); e) electrospun nanofibers’ diameter distribution; f) orientation analysis of SB and EHCS membrane nanofibers.

The specific stress metric strongly affected computed stress (Fig. 7a,b) and resistance to deformation (Fig. 7c,d), both for SB and EHCS. The material volume fraction (*f*) had the greatest impact that was amplified when considering its dependence on the imposed strain, as measured by micro-CT. In particular, volume fraction trend with strain appears realistic: in SB it slightly increased with strain because of bundle transversal contraption due to traction (Fig. 5a,b); in EHCS it decreased with strain because bundles-membrane spaces increased (Fig. 5c,d). The effect of strain is clear also comparing the differences between 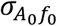 and 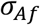 in Fig. 7a,b (*A*_0_ and *f*_0_ are the initial – out of micro-CT – section and volumetric fraction, while *A* and *f* are their strain-dependent update as measured in micro-CT). Assuming σ_*Af*_ as the most accurate metric, the results show that a first step forward an accurate evaluation of stress can be achieved by considering the initial volume fraction (*f*_*0*_) through Eq. 1. This metric does not require complex procedure and facilities. A better estimate can be achieved considering also cross-section area change with strain 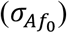. Though this change was evaluated through micro-CT scan in this study, a similar measure could be performed through macro-photos or videos during a standard monoaxial test and repeating the area evaluation at different strains, using the same procedure shown in Fig. 2a. Otherwise, a simplification of the experimental procedure is obtained by predicting the cross-section area change with strain with the first- or second-order models (Eq. 9 and Eq. 10). These models accurately replicated both the average and the specimen-specific area change (Fig. 8), considering equivalent Poisson’s moduli of 0.3 for SB and of 1.2 for EHCS, which are close to the same moduli computed from micro-CT scans. The use of metrics 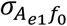 and 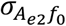does not require photos or videos during the test, and the subsequent image analyses.

**Figure 7.**
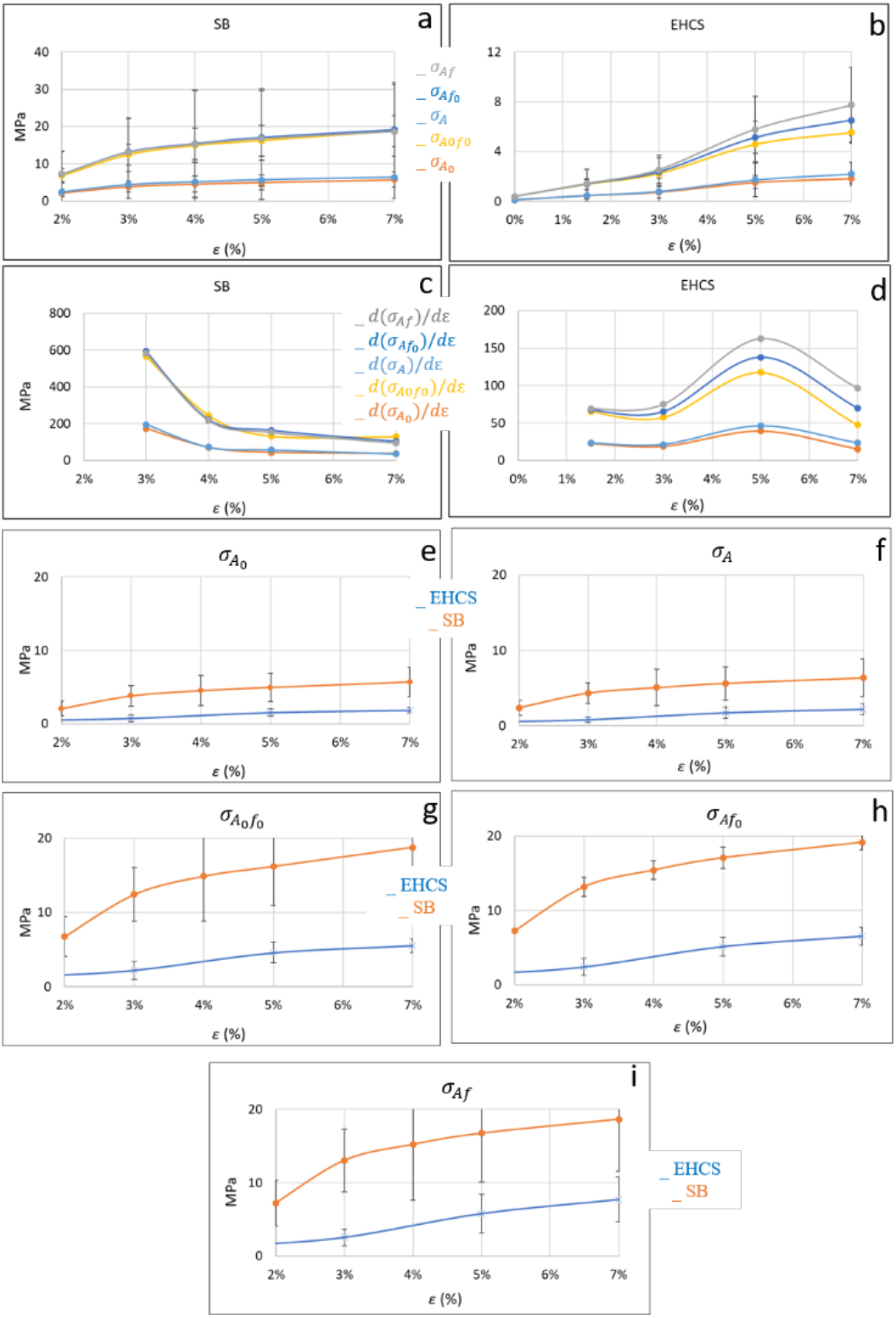
Curves of stress-strain (a, b, e-h; mean ± standard deviation) and of stress-strain gradients (c-d; mean) comparing the various samples (SB vs. EHCS) and the various stress calculations.

**Figure 8.**
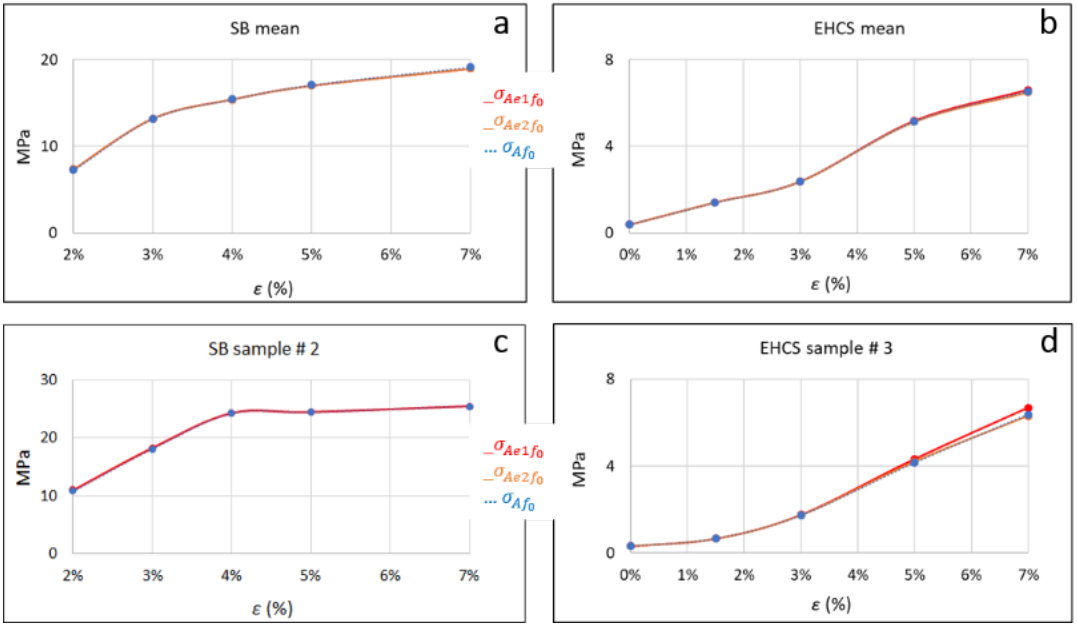
Curves of experimental 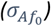 vs estimated 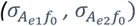 stress-strain (a,b: mean on SB,EHCS samples; c,d: specific samples SB #2,EHCS #3).

Differences between SB and EHCS stress-strain behavior were observed. The gap was also affected by the different metrics, with volume fraction still having the greatest impact and increasing the difference between the two (Fig. 7e-i). Whatever the stress metric, SB showed stress values higher than EHCS. Generally, also stress gradients were higher for SB at each strain level, though differences decreased at highest strains with uncrimping exhausted. Conversely, the gross shape of stress-strain curves, including their first derivatives, was not strongly affected by the chosen stress metric, both for SB and EHCS (Fig. 7a-d). Whatever the metric, indeed, (i) at 2-3% strain, EHCS still appeared in toe-region, showing higher non-linearity also because of evolving geometry (Fig. 9); (ii) non-linearity near yielding occurred around 3% strain for SB and around 5% for EHCS, though some EHCS samples reached higher values; (iii) stress derivative at 7% strain for EHCS reached values comparable to those in the toe-region (except for 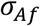, where it is higher), while for SB the same derivative is almost constant in the 5-7% strain range and always lower than the linear region; (iv) both SB and EHCS do not show failure until 7% strain.

**Figure 9.**
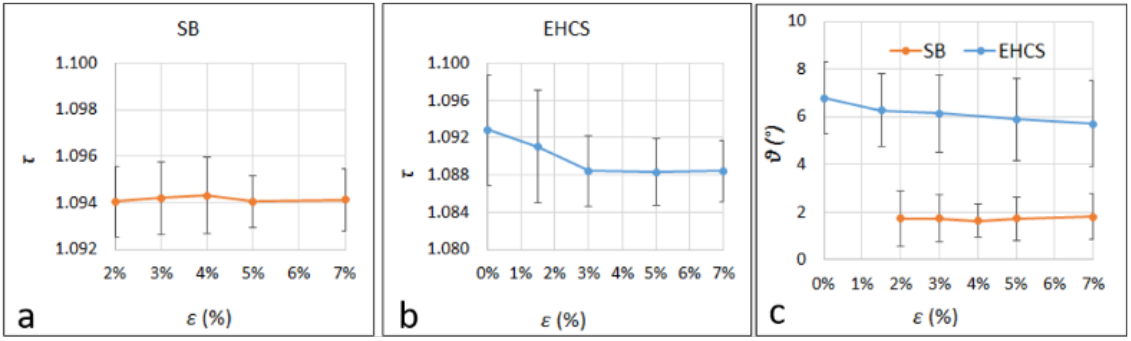
Bundles’ tortuosity-strain (a,b) and orientation-strain (c) graphs (mean ± standard deviation).

Since SB and EHCS share the same material and nanofibers geometry, differences can be ascribed to two factors, one at the micro-geometry and the other at the nano-geometry: (i) in EHCS, bundles showed a strain-dependent arrangement which appeared exhausted for SB, as described by orientation and tortuosity in Fig. 9 (decreasing for EHCS, stable for SB); (ii) the bio-inspired membrane enveloping the bundles of EHCS presents a random arrangement of nanofibers, while they are preferentially oriented along the loading axis in SB (Fig. 6f), therefore the membrane stress-shields the bundles, but offers less resistance to deformation. The membrane cross-section area was considered, though its contribution to strain resistance is lower than bundles. This also has an impact on elastic modulus (Fig. 7c,d, Tab. 1): its value at the end of the linear region (strain = 3% for SB, 5% for EHCS) computed using σ_*Af*_ is 583 MPa for SB and 163 MPa for EHCS on average. In addition, the load of fibers in EHCS is less uniform than in SB, due to the higher tortuosity and the composite nature of EHCS: while some fibers are just entering their linear region, others are in their non-linear or yielding region, so the actual resisting section is lower than the measured one. Previous DVC analyses confirmed a reduction in strain in some bundles with increasing load [20]. Moreover, inside EHCS, also the effect of crosslinking may contribute to the delay in the reorganization and alignment of bundles to reach their linear region. In fact, in EHCS the crosslinking of collagen can happen not only between the collagen molecules inside the single nanofiber, but also between nanofibers of different bundles or of bundles and membranes. This causes an increment of friction and a consequent delay in nanofibers/bundles alignment.

**Table 1.** Stress-strain gradients (mean± standard deviation) in the linear region (ε 3% for SB, 5% for EHCS) are assumed as elastic moduli of the tested samples and are statistically compared between sample types for each stress calculation method (SB vs. EHCS, ranksum test; * of different colours indicates significant difference between different pairs, p-value < 0.05) and between stress calculation methods for each sample type (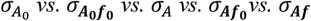, Kruskal-Wallis test; couple of letters indicates significant difference between pairs – small letters for SB, capital letters for EHCS –, p-value < 0.05).

| | $d(\sigma_{A_0})/d\varepsilon$ | $d(\sigma_{A_0f_0})/d\varepsilon$ | $d(\sigma_A)/d\varepsilon$ | $d(\sigma_{Af_0})/d\varepsilon$ | $d(\sigma_{Af})/d\varepsilon$ |
| --- | --- | --- | --- | --- | --- |
| **SB**<br>$\varepsilon$ 3% | 173±44 MPa<br>ad ae * | 565±93 MPa<br>* | 196±32 MPa<br>* | 594±97 MPa<br>ad * | 583±97 MPa<br>ae * |
| **EHCS**<br>$\varepsilon$ 5% | 39±26 MPa<br>AE * | 117±77 MPa<br>* | 47±30 MPa<br>* | 138±86 MPa<br>* | 163±107 MPa<br>AE * |

This study presents some limitations. Firstly, stress-strain behavior of the scaffolds can be considered reliable mainly in the elastic part, while yielding was strongly influenced by the clamping method and probably happened not in the analyzed ROI. However, the considerations on the impact of net vs apparent stress and of strain-dependent vs strain-independent physical quantities hold already at small strain (Fig. 10, Tab. 1). Secondly, micro-CT measures were affected by the low spatial resolution, in relation to the dimensional scale of the scaffolds’ architecture: nano-scale, e.g. nano-porosity, was lost, but also some micro-porosity due to partial volume effect and the voxel size adopted (i.e., a voxel was considered material when its signal was a combination of material and void contributions). For this reason, micro-CT porosity was used to update material volume fraction with evolving strain, but not to directly calculate it. In fact, while reference volume fraction – calculated by mass and density before testing – was about 0.3, micro-CT volume fraction at the minimum applied load resulted 0.8, unreliably high. However, as previously noted, its trend with strain was considered realistic. Third, load-cell sensitivity cannot guarantee an accurate initial no-strain starting point: the initial state has an initial strain that could be less accurate than other measures. This is the reason why the initial section (*A*_*0*_) updated for the strain-dependency – as evaluated by micro-CT – was taken as actual section (*A*) instead of the micro-CT measured section itself (*A’*). Using *A*_*0*_ as base parameter has also a practical reason: in presence of a valid model estimating section strain-dependency, as in this study, only loads must be read from a next traction test to obtain an estimate 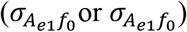 of reliable stress 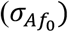, close to the actual one (σ_*Af*_) (no significant differences between the two metrics, e.g. see Table 1). Finally, to use *A’* instead of *A*, even if more accurate from the “actual” point of view, did not seem to significantly change the scenario (Supplementary Table S1 shows the same statistics of Table 1).

**Figure 10.**
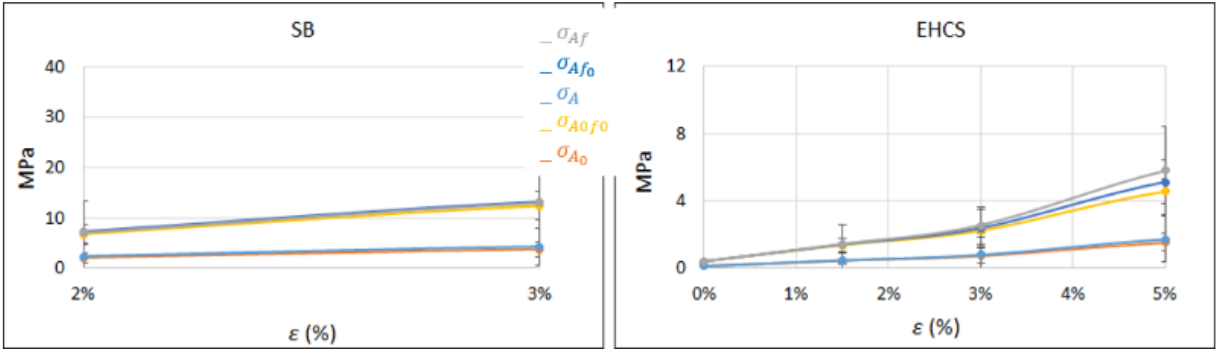
Curves of stress-strain (a SB, b EHCS; mean ± standard deviation) in the elastic region, comparing the various stress calculations.

The results of this study are both statistically and practically significant, having a potential functional/biological impact. Pereira et al. [35] also tested hybrid polymeric T/L scaffolds in tension by considering their solid cross section to calculate net stress. However, no previous study took explicitly in consideration the evolution of the scaffold’s morphology to reveal the actual stress-strain behavior. In the present study, statistical differences came out from the comparison between the elastic response of the different types of samples and between the different stress metrics (Tab. 1). As previously discussed, EHCS showed a different stress-strain behavior with respect to SB, being progressively and non-uniformly stretched, resulting in extended non-linear region at low (toe region) and high (yielding region) strain. This has a functional impact: when implanted in substitution or support of a T/L, EHCS resist to deformation in a biomimetic way. Moreover, there is a potential biological impact: thanks to the membrane, cells face a lower substrate stiffness in EHCS with respect to SB [27][13]. The general importance of this factor on cellular mechano-transduction is well known [36]. Specifically to the tested scaffolds, in terms of stiffness SB and EHCS showed stress-strain gradient close respectively to the first and second substrate group presented in the study of Islam et al. [37], which found out how the higher substrate stiffness increased tenogenic differentiation. In addition, only considering net material and geometry dependence on strain, the stress calculation actually differed from the standard apparent stress (Tab. 1). This has a functional impact: the composite structures tested here show an apparent stress distant from the native T/L mechanical response, while the actual stress falls in the range of tissues with similar dimensions, making the studied scaffolds quite promising [2][15]. Moreover, there is a potential biological impact: σ_*Af*_ gives a measure of the actual substrate stiffness and geometry that cells will face *in vivo*. Our measures indicate for EHCS a σ_*Af*_ stress-strain gradient of 163 MPa (Tab. 1), close to the stiffest – and most effective in terms of tenogenic differentiation – substrate studied by Islam et al. [37].

## Conclusion

Considering strain-dependent morphology is crucial for describing the actual mechanical behavior – e.g. stress-strain relation – of micro/nano-porous scaffolds. This study showed it by a micro-CT *in situ* test on novel hierarchical composite tendon/ligament scaffolds. Therefore, it suggests using strain-dependent net stress as reference metric for scaffold development.

## Acknowledgments

The Proof Of Concept Grant of the University of Bologna and the Horizon Europe Marie Skłodowska Curie Postdoctoral Fellowship 3NTHESES (Grant No. 101061826) are greatly acknowledged. This work was done while A. Sensini was within the Department of Industrial Engineering, University of Bologna, Italy. Type I collagen was kindly provided by Kensey Nash Corporation d/b/a DSM Biomedical (Exton, USA).

## Author contributions

Gregorio Marchiori: Writing – review & editing, Writing – original draft, Methodology, Investigation, Formal analysis, Data curation, Visualization, Conceptualization. Nicola Sancisi: Writing – review & editing, Writing – original draft, Methodology, Investigation, Formal analysis, Conceptualization. Gianluca Tozzi: Writing – review & editing, Writing – original draft, Methodology, Conceptualization. Massimiliano Zingales: Formal analysis, Writing – review & editing. Gaia Prezioso: Formal analysis, Writing – review & editing. Andrea Visani: Writing – review & editing, Supervision. Andrea Zucchelli: Writing – review & editing, Writing – original draft, Supervision, Methodology, Funding acquisition, Project administration, Conceptualization. Alberto Sensini: Writing – review & editing, Writing – original draft, Visualization, Project administration, Methodology, Investigation, Funding acquisition, Formal analysis, Data curation, Conceptualization.

## Funding

This work was supported by The Proof Of Concept Grant of the University of Bologna and by the Italian Ministry of Health [grant number 5×1000 2022-23685320].

## Supplementary Material

**Video:** Video S1, Video S2

**Table:** Table S1

**Video S1.**
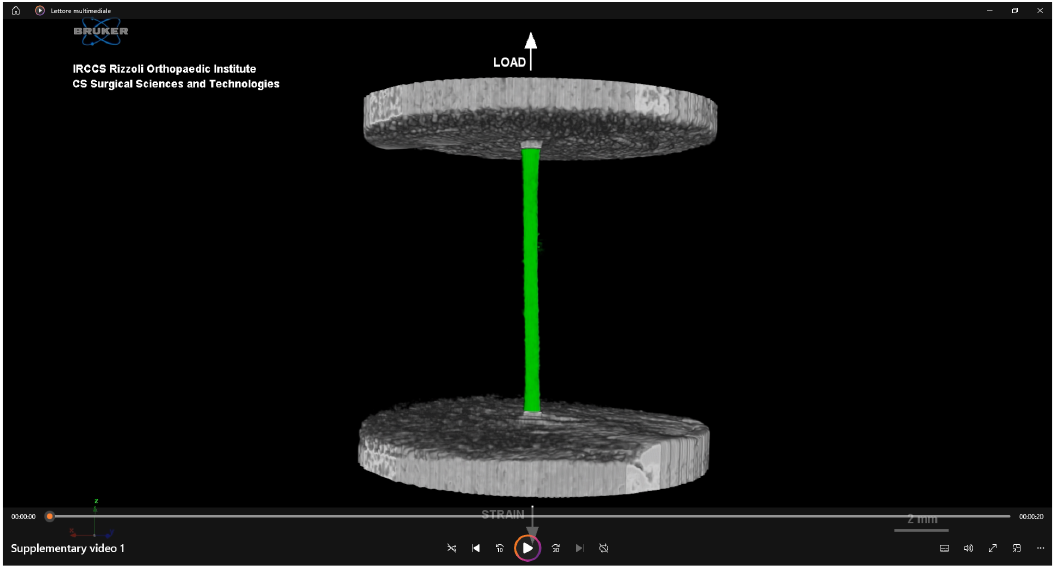
3D rendering video created by CT-Vox software of the micro-CT in situ test and analysis of an electrospun tendon/ligament bundle scaffold.

**Video S2.**
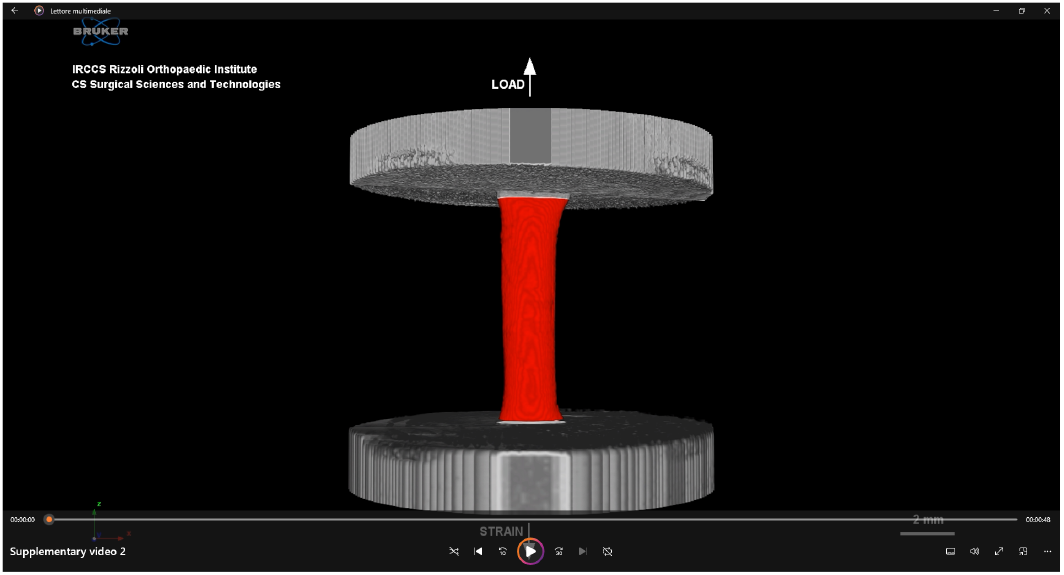
3D rendering video created by CT-Vox software of the micro-CT in situ test and analysis of an electrospun tendon/ligament composite hierarchical scaffold.

**Table S1.**
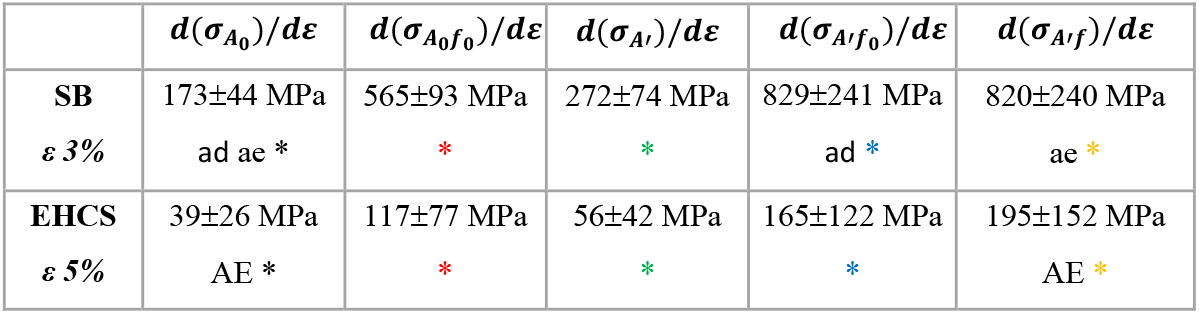
Stress-strain gradients (mean± standard deviation) in the linear region (ε 3% for SB, 5% for EHCS) are assumed as elastic moduli of the tested samples and are statistically compared between sample types for each stress calculation method (SB vs. EHCS, ranksum test; * of different colours indicates significant difference between different pairs, p-value < 0.05) and between stress calculation methods for each sample type (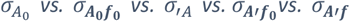, Kruskal-Wallis test; couple of letters indicates significant difference between pairs – small letters for SB, capital letters for EHCS –, p-value < 0.05).

## Notes

### Competing Interest Statement

Andrea Zucchelli and Alberto Sensini have patent ##EP3638328 issued to University of Bologna.The other authors declare that they have no known competing financial interests or personal relationships that could have appeared to influence the work reported in this paper.

